# Sex, androgens and regulation of pulmonary AR, TMPRSS2 and ACE2

**DOI:** 10.1101/2020.04.21.051201

**Authors:** Mehdi Baratchian, Jeffrey M. McManus, Mike Berk, Fumihiko Nakamura, Sanjay Mukhopadhyay, Weiling Xu, Serpil Erzurum, Judy Drazba, John Peterson, Eric A. Klein, Ben Gaston, Nima Sharifi

## Abstract

The sex discordance in COVID-19 outcomes has been widely recognized, with males generally faring worse than females and a potential link to sex steroids. A plausible mechanism is androgen-induced expression of TMPRSS2 and/or ACE2 in pulmonary tissues that may increase susceptibility or severity in males. This hypothesis is the subject of several clinical trials of anti-androgen therapies around the world. Here, we investigated the sex-associated TMPRSS2 and ACE2 expression in human and mouse lungs and interrogated the possibility of pharmacologic modification of their expression with anti-androgens. We found no evidence for increased *TMPRSS2* expression in the lungs of males compared to females in humans or mice. Furthermore, in male mice, treatment with the androgen receptor antagonist enzalutamide did not decrease pulmonary TMPRSS2. On the other hand, ACE2 and AR expression was sexually dimorphic and higher in males than females. ACE2 was moderately suppressible with enzalutamide therapy. Our work suggests that sex differences in COVID-19 outcomes attributable to viral entry are independent of TMPRSS2. Modest changes in ACE2 could account for some of the sex discordance.

## Introduction

The December 2019 outbreak caused by the severe acute respiratory syndrome coronavirus 2 (SARS-CoV-2) in Wuhan, China, has led to the coronavirus disease 2019 (COVID-19) pandemic (1, 2). The highly contagious nature of the disease as well as the lack of vaccines or clinically approved treatments has caused a worldwide public health emergency. Therefore, improved and timely understanding of the human susceptibilities to SARS-CoV-2 will prove invaluable in controlling the pandemic and treatment of those affected.

SARS-CoV-2 enters cells using two host cellular proteins: angiotensin converting enzyme-2 (ACE2) and transmembrane serine protease 2 (TMPRSS2). The virus first employs ACE2 as a cell entry protein, followed by TMPRSS2-mediated proteolytic processing of the SARS-2 spike protein, further facilitating viral entry (3). Targeting the activity or expression of both factors by a plethora of approaches has been proposed as potential treatment (3–5). *TMPRSS2* is also a widely studied androgen-regulated gene in prostate tissue, contributing to prostate cancer pathogenesis by way of aberrantly driving oncogene expression. Approximately half of all prostate cancers harbor a fusion that juxtaposes a *TMPRSS2* transcriptional regulatory element, which is stimulated by potent androgens and the androgen receptor (AR), in front of an ERG oncogene. (6). The end result is AR stimulation of oncogene expression which promotes growth of prostate cancer. However, two population-based studies of men undergoing hormonal therapy for prostate cancer have yielded differing results on a possible protective effect of androgen suppression on risk of COVID-19 (7, 8).

Androgen regulation of TMPRSS2 raises the possibility that the physiological roles of androgens may, at least partially, account for the sex-specific clinical outcomes (9, 10). Utilizing a high-throughput drug screening strategy, a recent study found that ACE2 levels in human alveolar epithelial cells can be downregulated by 5α-reductase inhibitors, suggesting an androgen-driven mode of expression (11). Furthermore, due to its androgen-regulated nature in the prostate and its essential role in SARS-CoV-2 etiology, TMPRSS2 expression has been postulated to follow a similar pattern of regulation in pulmonary cells by the potent androgens testosterone and dihydrotestosterone (12). If this link proves correct, it could pave the path to novel strategies, including re-purposing of FDA-approved potent androgen synthesis inhibitors or AR antagonists, such as enzalutamide (Enz) and apalutamide, for the treatment of COVID-19. These strategies are the subject of several clinical trials (e.g., NCT04374279, NCT04475601, NCT04509999, NCT04397718) (5, 13).

Here, we show that the expression of pulmonary AR and ACE2 follows a sex-discordant pattern with males expressing considerably higher levels of protein than females. In humans, there is no difference in ACE2 expression between non-smoking men and women, while in contrast, ACE2 expression is significantly higher in the lungs of male smokers. We provide *in vivo* evidence that neither mRNA nor protein levels of TMPRSS2 vary by sex or treatment with the potent AR-antagonist Enz. ACE2 expression however is modestly modifiable by anti-AR treatment and may to some extent explain the sex disparities in susceptibility to SARS-CoV-2.

## Results

### Sexually dimorphic AR expression and ACE2 dimorphism in smokers

Certain pulmonary disease outcomes, including asthma, are sex steroid-associated (14). Considering the poorer clinical outcome of COVID-19 in men, underlying androgen-related causes are suspected but not presently known. The SARS-CoV-2 co-receptor TMPRSS2 harbors an AR-responsive enhancer that is induced by androgens in prostate tissue (15), raising the possibility of a similar mode of regulation in the respiratory system. We first asked whether, similar to TMPRSS2, ACE2 was also regulated by AR signaling in LNCaP, which is AR expressing and androgen responsive. Indeed, both mRNA and protein expression of ACE2 were strongly induced by the synthetic androgen R1881 and suppressed by Enz-mediated AR blockade (**Fig. 1A** and **1B**). Moreover, ChIP-seq analysis of AR cistrome revealed multiple AR-binding sites upstream of the ACE2 region that were lost upon Enz treatment (**Fig. 1C**). These findings collectively indicate that ACE2 is indeed an androgen-driven gene in prostate cells.

**Figure 1.**
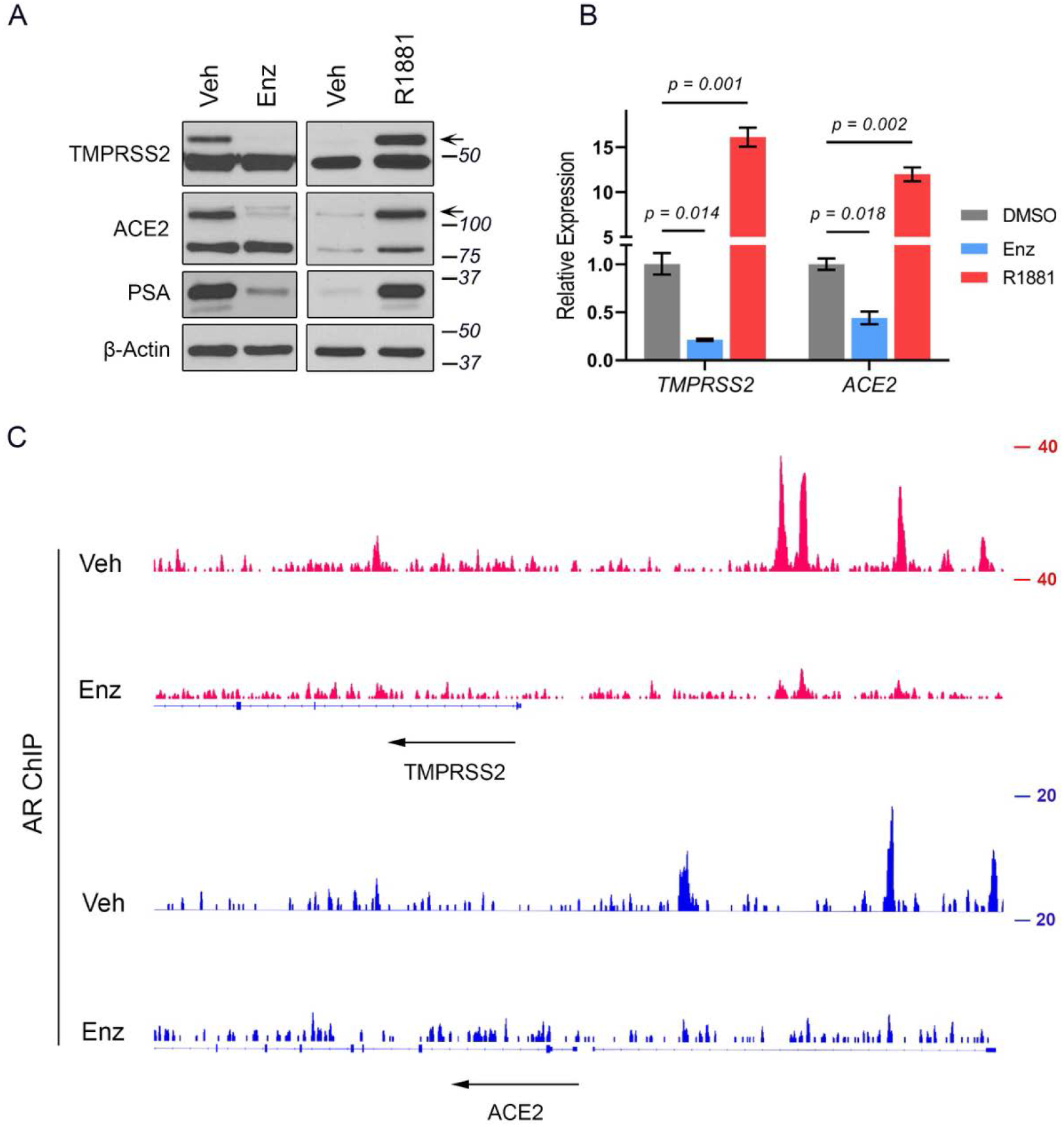
ACE2 is an androgen-regulated gene in prostate cells. A) Immunoblots and B) RT-qPCR analysis of TMPRSS2 and ACE2 expression in LNCaP cells treated with Enz (10 μM) for 14 days or stimulated with R1881 (5 nM) for 48 hours. Vehicle (Veh) used for Enz and R1881 were DMSO and ethanol, respectively. Results (mean±SD) are representative of three biological repeats, performed in triplicate. p values were determined using one-way ANOVA. Arrows indicate the location of specific bands. C) ChIP-seq track examples of AR occupancy within TMPRSS2 and ACE2 gene regions, in LNCaP cells treated with Veh (DMSO) or Enz (5 μM) for 14 days.

We next sought to investigate whether male sex was associated with higher expression of *ACE2, TMPRSS2* or *AR* in human lung. To this end, we acquired the publicly available expression datasets in non-cancerous lung and associated respiratory tissues from the Genomic Expression Omnibus (GEO). Across all tissue-type comparisons, we found no evidence for elevated *ACE2, TMPRSS2* or *AR* mRNA expression in males compared with females (**Fig. 2**). Next, we performed immunohistochemical (IHC) analyses to explore the possibility of a sex-specific pattern of protein expression within distinct pulmonary cell types. The IHC-stained mouse and human lung specimens were assessed by an expert pulmonary pathologist (SM) and semi-quantified for protein expression using the H-score, a common and well-known semiquantitative scoring system that takes into consideration staining intensity as well as the percentage of cells staining positively (16–18). H-scores can range from 0 (no expression) to 300 (3+ staining intensity in 100% of cells).

**Figure 2.**
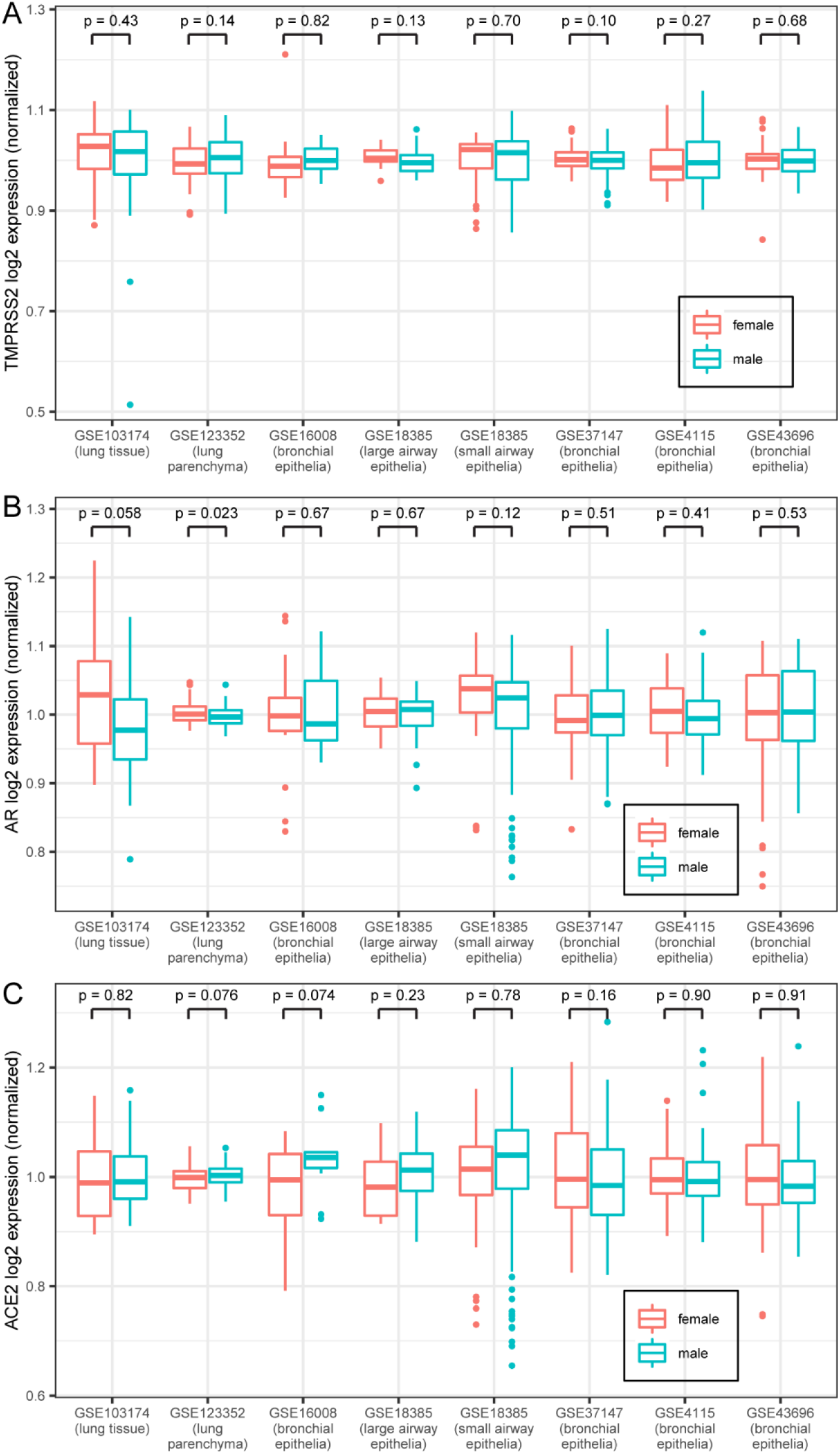
TMPRSS2, AR, and ACE2 transcript expression in human lung are similar in males and females. (A) Box plot of TMPRSS2 expression (normalized so that the mean within each data set equals 1) from the publicly available Gene Expression Omnibus (GEO) data sets. Center lines indicate median values, edges of boxes indicate first and third quartile values, whiskers indicate largest and smallest values extending no more than 1.5 * inter-quartile range from edges of boxes, and dots indicate outlier values. P-values from t-tests are shown for female vs. male comparison within each data set. N for each data set: GSE103174 = 22 female/31 male; GSE123352 = 81 female/95 male; GSE16008 = 15 female/11 male; GSE18385 large airway = 16 female/36 male; GSE18385 small airway = 35 female/74 male; GSE37147 = 103 female/135 male; GSE4115 = 41 female/122 male; GSE43696 = 74 female/34 male. Data sets 18385 and 4115 contained multiple TMPRSS2 reference sequences; none of the individual sequences had female vs. male expression differences with p < 0.05, and the expression values of the different sequences were added together for this plot. (B) Box plot of AR expression from the same data sets as in (A). (C) Box plot of ACE2 expression from the same data sets as in (A).

Pulmonary TMPRSS2, predominantly expressed in airways (bronchial or bronchiolar epithelial cells), was neither associated with sex in humans (**Fig. 3A**) nor in mice (**Fig. 3D**). Furthermore, smoking did not appear to affect TMPRSS2 expression (**Fig. 3A**). In human lungs, alveolar epithelial cells also stained positive for TMPRSS2; expression was unchanged by smoking status or sex.

**Figure 3.**
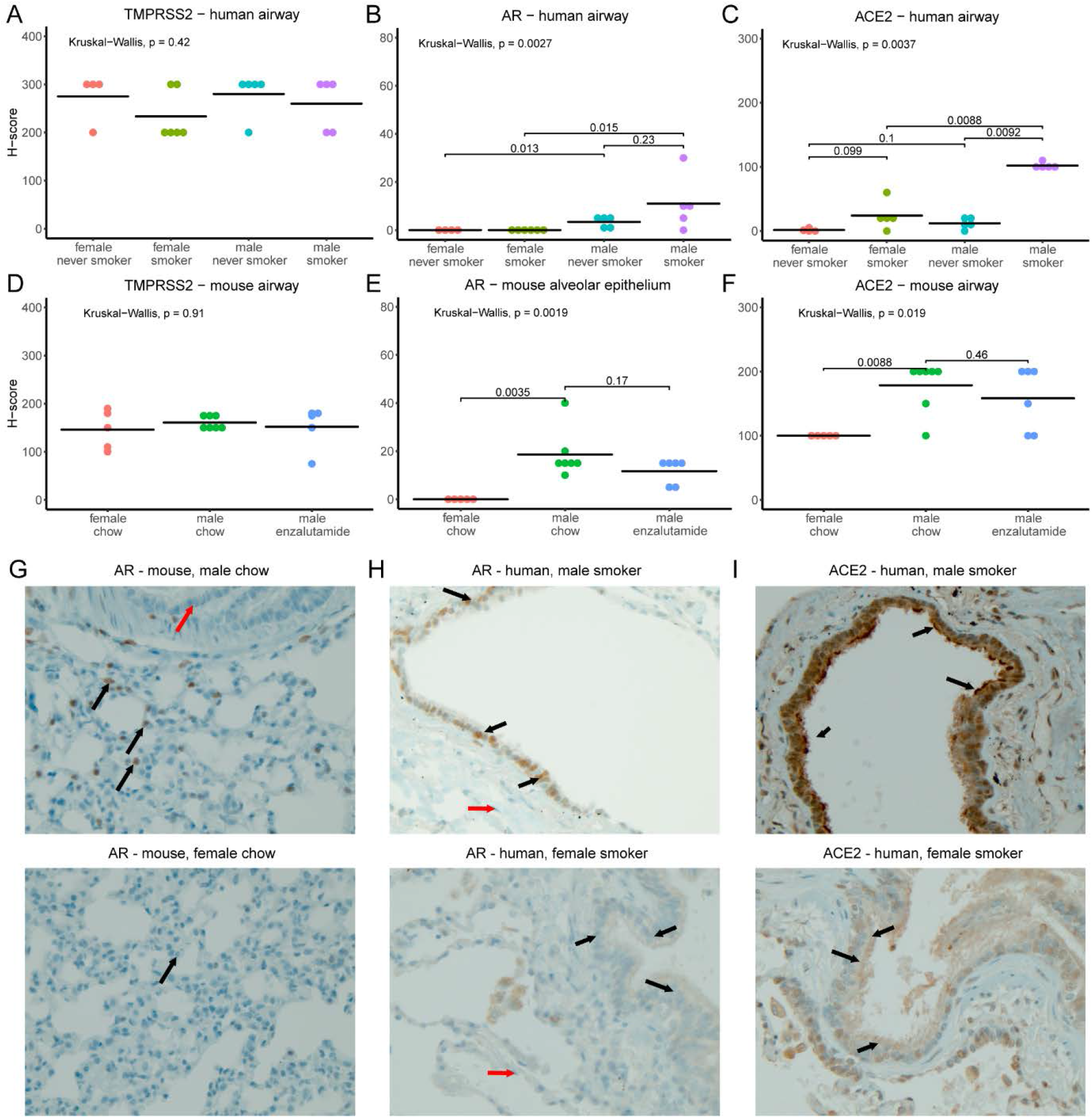
AR and ACE2, but not TMPRSS2, display sex-discordant patterns of immunohistochemical staining in human and mouse lung. (A-F) Plots of H-scores from staining for TMPRSS2 (A, D), AR (B, E), and ACE2 (C, F) in human (A-C) and mouse (D-F) lung samples. Note that for AR in mouse, alveolar epithelial scores are shown, as airway scores were zero for all samples; for all other panels, airway scores are shown. Humans were smokers or never-smokers as indicated. Male mice were given control chow or enzalutamide as indicated whereas female mice were given control chow. Plots show each individual data point and horizontal lines for mean values. P-values from Kruskal-Wallis tests are shown, and for plots in which Kruskal-Wallis tests suggested differences between groups, p-values from post hoc Wilcoxon tests are shown. (G-I) Representative images (400x magnification) illustrating sex differences in staining. (G) AR staining in control male (top) and female (bottom) mice. Black arrows indicate alveolar epithelial cells with positive nuclear staining in male and negative in female. Red arrow indicates lack of staining in bronchiolar epithelial (airway) cells. (H) AR staining in human male smoker (top) and female smoker (bottom). Black arrows indicate airway epithelial cells with positive nuclear staining in male and absence of staining in female. Red arrows indicate alveolar epithelial cells negative in both. (I) ACE2 staining in human male smoker (top) and female smoker (bottom). Black arrows indicate positive apical membrane staining in male airway epithelial cells and absence of staining in female airway epithelial cells.

In contrast, we discovered that AR and ACE2 demonstrate sex-discordant expression; AR showed limited expression in airway cells of human males which was absent in human females (**Fig. 3B**). Smoking further elevated AR levels to a modest degree in men but the difference did not reach statistical significance (**Fig. 3B** and **3H**). Consistent with the overall pattern observed in humans, lungs of male mice expressed markedly higher levels of AR protein than females (**Fig. 3E** and **4D**). Alveolar pneumocytes (epithelial cells) were the dominant AR-expressing population in mouse lungs (**Fig. 3G**). Since the AR transcript levels did not differ by sex (**Fig. 2B** and **4C**), we speculated that the high concentrations of circulating androgens in males may stabilize pulmonary AR expression (19).

**Figure 4.**
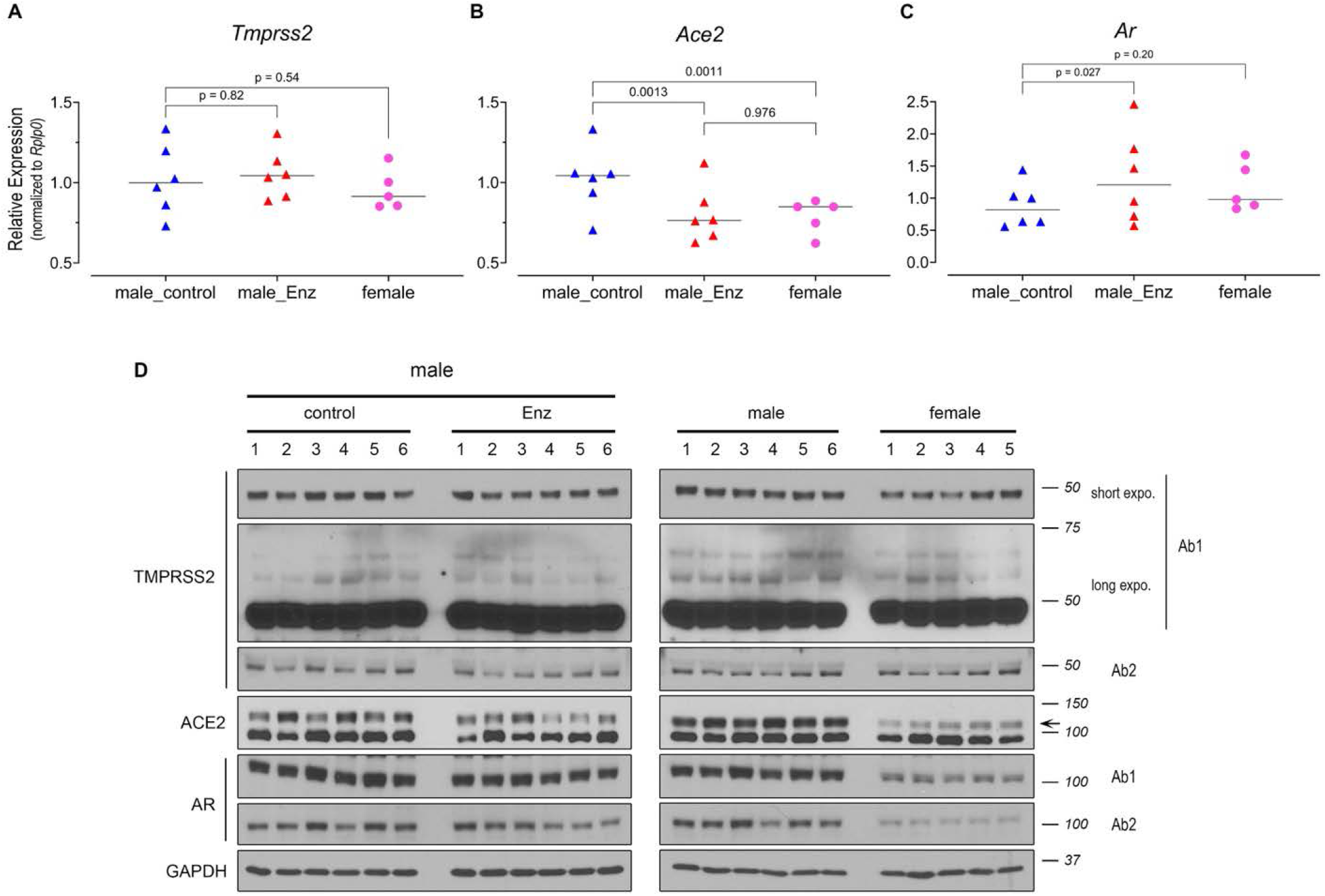
AR and ACE2 protein but not TMPRSS2 present a pattern of sex discordance in mouse lung. **(A-C)** Transcript levels of *Tmprss2* (A), *Ace2*, (B) *Ar* (C) in female (n=5) and male NSG mice treated with control (n=6) or Enz diet (n=6). Gene expression was assessed in triplicate and normalized to *Rplp0* levels. The statistical differences were calculated using one-way ANOVA with Tukey’s post hoc test. Results are shown as mean ± s.d. (n = 3 technical repeats). **(D)** Immunoblots showing the expression of pulmonary TMPRSS2, ACE2 and AR in male mice fed with Enz or control chow for 11 days, and in male vs female mice. Results are representative of 4 technical repeats. TMPRSS2 (Ab1: ab92323; Ab2: 14437-1-AP) and AR (Ab1:PG-21 and Ab2: N-20). Arrow indicates the location of ACE2.

In the case of ACE2, we detected no significant difference between non-smoker men and women. Nevertheless, its expression was elevated in the lungs of male smokers compared with female smokers (**Fig. 3C** and **3I**). Thus, we observed sexually dimorphic ACE2 expression in smokers only. In mice, the sex disparity was readily detectable, with males expressing significantly higher ACE2 protein in airways than females (**Fig. 3F**). These data, together, suggest that pulmonary ACE2 and AR expression is sexually dimorphic.

### TMPRSS2 expression is unaffected and ACE2 modestly suppressed by potent AR blockade

To further explore the possibility of inhibiting TMPRSS2 expression by means of AR blockade, we harvested lungs from female and male mice treated with control diet or Enz for > 10 days and analyzed them for protein and mRNA expression. The animal studies performed on bulk lungs yielded results consistent with the human expression analysis: i.e., there were no sex-specific TMPRSS2 changes. Additionally, Enz treatment failed to detectably downregulate TMPRSS2 (**Fig. 4A** and **4D**). It has been previously noted that, due to glycosylation, full-length TMPRSS2 may be detected by SDS-PAGE at a higher molecular weight of approximately 70 kDa (20). In immunoblots performed with maximum-sensitivity substrate and extended exposures, we did not detect distinguishable bands for TMPRSS2 above 50 kDa (Fig. 4D). Given the unchanged AR protein levels (despite a modest but statistically significant mRNA increase) in Enz-treated males, we infer that TMPRSS2 is not regulated by AR in the lung. Nevertheless, high resolution techniques such as single cell-sequencing or -proteomics may be employed to explore otherwise undetectable alterations of expression in minority cell populations.

In keeping with human data, we found no evidence of sex-specific changes in *AR* transcript levels in mice (**Fig. 4C**). The lungs of male mice, however, expressed significantly higher amounts of AR protein compared with females (**Fig. 4D**). As mentioned previously, this may be due to AR stabilizing effects of abundant circulating androgens in males. Similar to AR, ACE2 expression also showed clear sexual dimorphism, with males expressing substantially higher levels of protein (**Fig. 4D**) and modestly but significantly higher amounts of mRNA (**Fig. 4B**) in bulk lungs compared with females. Treatment with Enz lowered the transcript levels in males down to female levels (**Fig. 4B**) and also partially reduced protein quantities (**Fig. 4D**) as evidenced by immunoblotting. The modest Enz-mediated suppression of ACE2 was not clearly captured in our IHC analyses. This could be explained by possible marginal changes of ACE2 within pulmonary cells that may fall below the IHC detection limit, however, accumulatively can be observed through total protein detection methods performed on bulk tissue. Overall, these data indicate sex-discordant AR and ACE2 regulation, and a potential androgen-regulated mode of expression for pulmonary ACE2. Therefore, androgen-AR-mediated mechanisms could explain sex-specific differences in COVID-19 outcomes by TMPRSS2-independent mechanisms.

### *ACE2* and *TMPRSS2* mRNA expression increases in current smokers whereas in former smokers, expression returns to levels found in never-smokers

In addition to male sex, smoking is a risk factor for COVID-19 susceptibility and poor clinical outcomes (21). One recent study of 1,099 COVID-19-positive patients reported a more than two-fold increased risk for intensive care unit admission and death in smokers as compared with non-smokers (22). We identified human expression GEO datasets of bronchial/airway epithelial cells containing subject smoking status and asked whether smoking is associated with *TMPRSS2* expression. Our analysis indicated a consistent pattern whereby expression of both *TMPRSS2* (**Fig. 5A**) and the primary SARS-CoV-2 receptor *ACE2* (**Fig. 5B**) was modestly but significantly increased in smokers compared with non-smokers. Interestingly, the levels were downregulated to never-smoker levels in former smokers. The results of our analysis are in keeping with several recent reports on ACE2 and smoking (23–25). Although the p-values range widely across different data sets, almost all show changes in a consistently increased direction for current smokers including multiple data sets with small p-values (**Fig. 5A** and **5B**). Finally, for both *TMPRSS2* (**Fig. 5C**) and *ACE2* (**Fig. 5D**), there was no correlation between smoking pack-years and mRNA expression in either current or former smokers, suggesting that the change does not build up over time but is instead a rapid process akin to a switch.

**Figure 5.**
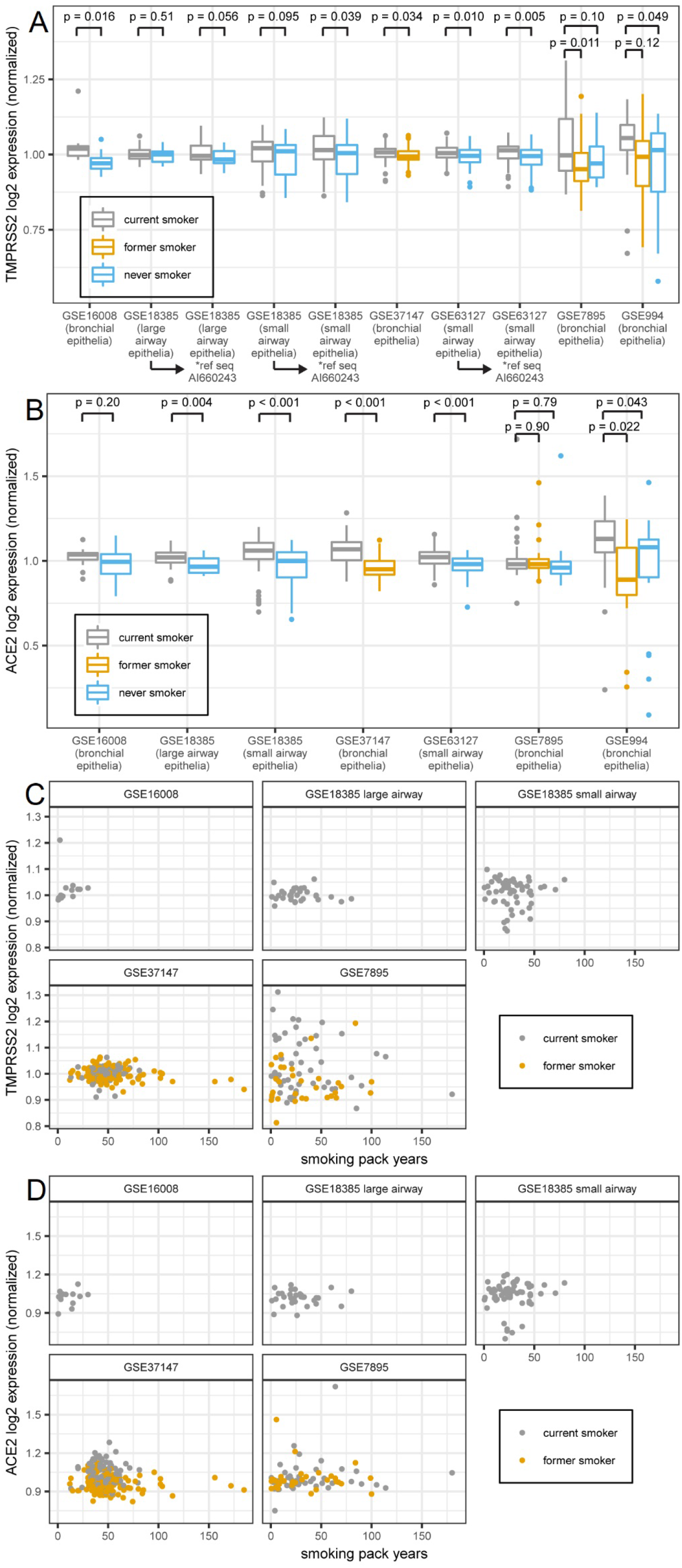
Expression of both TMPRSS2 and ACE2 transcript in human bronchial epithelia increases in current smokers compared to both former and never smokers and the increases do not depend on smoking pack years. (A) Box plot of TMPRSS2 expression (normalized; mean within each data set equals 1) from GEO data sets. N for each data set: GSE16008 = 13 current/13 never smoker; GSE18385 large airway = 32 current/20 never smoker; GSE18385 small airway = 58 current/51 never smoker; GSE37147 = 99 current/139 former smoker; GSE61327 = 112 current/71 never smoker; GSE7895 = 52 current/31 former/21 never smoker; GSE994 = 34 current/18 former/23 never smoker. Data sets 18385, 63127, 7895, and 994 contained multiple TMPRSS2 reference sequences whose expression values were summed. In all data sets containing four reference sequences (18385 large and small airway and 63127), the difference between groups was largest for sequence AI660243, so expression using that sequence alone is also shown. For data sets with two groups, p-values were obtained from t-tests. For data sets with three groups, Tukey HSD p-values were obtained after one-way ANOVA (ANOVA p-values: GSE7895 0.009, GSE994 0.033). (B) Box plot of ACE2 expression from the same data sets as in (A). Data sets 18385 and 63127 contained multiple ACE2 reference sequences whose expression values were summed. For data sets with two vs. three groups, p-values were obtained as in (A) (ANOVA p-values for three group sets: GSE7895 0.82, GSE994 0.010). (C) Scatter plots of normalized TMPRSS2 expression vs. smoking pack years for current and former smokers in data sets containing pack year data. GSE37147 (former smokers) had adjusted R^2^ = 0.03 and p = 0.03; no other linear regression had p < 0.1. (D) Scatter plots of normalized ACE2 expression vs. smoking pack years. No linear regression had p < 0.1.

## Discussion

Sex-associated clinical outcomes have been long observed in a variety of infectious and inflammatory conditions. Sex steroids (i.e., androgens and estrogens) are possible mediators of these biologic differences. For COVID-19, potential androgen-mediated biologic differences include 1) SARS-CoV2 cellular receptor regulation and 2) immune modulation (5, 13). Our study addresses the first of these possibilities.

In both humans and mice, AR protein is clearly more highly expressed in the lungs of males compared with females. Detection of AR protein in male compared with female lung might be surprising in the absence of a detectable difference in AR transcript. This might be explained by stabilization of AR protein in the presence of androgens, which is recognized to occur in prostatic tissues (19). To our knowledge, there are no prior reports of this sexually dimorphic pulmonary AR expression. The specific presence of AR protein in male lungs raises the question of its transcriptional program – namely expression of TMPRSS2 and ACE2.

We find no evidence for androgen regulation of *TMPRSS2* in pulmonary tissues. This evidence includes 1) the absence of any *TMPRSS2* increase in male compared with female human lung 2) no *TMPRSS2* increase in male compared with female mouse lung 3) no evidence for *TMPRSS2* suppression with next-generation AR antagonist treatment. Our observations are in agreement with a new study that provides evidence for the absence of AR-binding and open chromatin state within the TMPRSS2 locus in lung as compared with prostate cells (26). We do find evidence for an increase in TMPRSS2 transcript but not protein expression with smoking. This difference in findings for transcript vs. protein expression might be attributable to differences in the biospecimens sampled across studies or differential expression in cellular subtypes.

In contrast to TMPRSS2, we do find evidence for sexually dimorphic ACE2 expression. Specifically, protein expression is higher in the lungs of male smokers compared with female smokers and in male mice compared with female mice. In terms of pharmacologic intervention, suppression of ACE2 expression with a potent AR antagonist in the lungs of male mice is significant albeit modest. Whether this apparent magnitude of androgen regulation of ACE2 expression is meaningful for SARS-CoV-2 infection or COVID-19 severity is unclear. Nevertheless, this will be tested in several ongoing clinical trials (including ClinicalTrials.gov NCT04475601, NCT04397718, NCT045009999 and NCT04374279).

Sexually dimorphic AR expression in an organ not associated with sexual differentiation – the male lung – raises the question of function. This is reminiscent of sexually dimorphic AR protein expression in the male human kidney, in which AR function includes regulating glucocorticoid metabolism and downstream steroid receptor activity (27). Whether this physiology also occurs in the lung has yet to be determined. This may have implications for regulation of inflammatory processes including asthma.

In conclusion, we find no evidence for androgen regulation of *TMPRSS2* in the male lung. Therefore, *TMPRSS2* regulation in the lung appears to fundamentally differ from clear androgen-dependent regulation in prostatic tissues. In contrast, there is a sex discordance in AR and ACE2 expression in lungs of mice and humans. In humans, elevated ACE2 expression is apparent in the lungs of male smokers compared with female smokers. Pulmonary TMPRSS2 regulation appears not to account for the sex-discordance in COVID-19 clinical outcomes. In contrast, ACE2 expression is elevated in smokers and particularly in males. The magnitude of ACE2 suppression with enzalutamide in mouse is modest. The ultimate effects of anti-androgens on human pulmonary ACE2 expression and COVID-19 outcomes are not yet known. Smokers could partly mitigate their increased risk by quitting smoking.

## Materials and methods

### Mice, treatments and lung harvest

A cohort of adult NSG mice (> 6 weeks old) were obtained from Cleveland Clinic Biological Resources Unit. The male mice were arbitrarily divided between two groups receiving control chow or Enz diet 62.5 mg/kg. Following 11 days on diet, the mice were sacrificed using a lethal dose of Nembutal followed by cardiac puncture. Once sacrificed, the abdominal and thoracic cavities of the mice were opened, the inferior vena cava was cut, and the lungs were gently perfused with warm saline via the right ventricle. Next, the lungs were removed, and the individual lobes were either fixed in 10% formalin, or embedded in paraffin, or snap frozen for subsequent RNA or protein analysis.

### Protein and mRNA expression analysis

Approximately 40–50 mg freshly frozen lung was added to soft tissue homogenizing CK14 tubes (Betin Technologies) with 200 ml RIPA buffer containing HALT protease and phosphate inhibitor cocktail. Lung tissues were then homogenized with a homogenizer (Minilys, Betin Technologies) three times (60 s each time) at the highest speed, with 5-10 minute intervals on ice to cool lysates. The lysates were then centrifuged for 15 min at 16,000 x g and the supernatants were collected for immunoblot analysis with antibodies for TMPRSS2 (Abcam: ab92323 and Proteintech: 14437-1-AP), AR (EMD Millipore: PG-21 and Santa Cruz Biotechnology: N-20), PSA (Cell signaling: D6B1) and GAPDH (D16H11).

Total RNA was harvested by homogenizing 25 mg lung tissue in 350 μl RLT buffer (RNeasy kit, Qiagen) following the manufacturer’s instructions. cDNA synthesis were then carried out with the iScript cDNA Synthesis Kit (Bio-Rad). Quantitative PCR (qPCR) analysis was conducted in triplicate in an ABI 7500 Real-Time PCR machine (Applied Biosystems) using iTaq Fast SYBR Green Supermix with ROX (Bio-Rad) with the following primer sets: Tmprss2 - forward gtcatccacacacatcccaagtc; reverse tcccagaacctccaaagcaaga; Ace2 - forward actatgaagcagagggagcagatg; reverse ggctgatgtaggaagggtaggtat; Ar - forward ggcagcagtgaagcaggtag; reverse cggacagagccagcggaa; Rplp0 -forward gacctccttcttccaggctttg; reverse ctcccaccttgtctccagtcttta; TMPRSS2 - forward atcggtgtgttcgcctctacg; reverse atccgctgtcatccactattcctt; ACE2 - forward ggaggatgtgcgagtggcta; reverse taggctgttgtcattcagacgg; RPLP0 – forward attacaccttcccacttgctg; reverse actcttccttggcttcaacctta.

### ChIP-seq

ChIP-Seq analysis was performed in LNCaP cells treated with vehicle (DMSO) or Enz 10 μM. Briefly, 106 cells were cross-linked using 1% formaldehyde (reconstituted in 1X PBS) at room temperature for 10 min, followed by quenching with Glycine (final concentration 125 mM) and further incubation at room temperature for 5 min. Fixed cells were lysed in SDS buffer (50 mM Tris-HCl pH 8, 1% SDS, 10 mM EDTA), and sonicated at 4 degrees using Bioruptor sonicator (Diagenode) for 20 cycles on high setting: 10” on/10” off. The lysates were then immunoprecipitated in ChIP Dilution Buffer (20 mM Tris-HCl pH 8, 1% Triton-X 100, 2 mM EDTA, 150 mM NaCl + PIC), using 3 μg chromatin, 10 μl of anti-AR (Millipore, EMD Millipore: PG-21), and 50 μl of blocked A/G beads. Recovered DNA was used to prepare libraries using the Illumina Nextera library prep method, subsequently sequenced on a NextSeq 500 and analyzed using the ChiLin analytical pipeline. Finally, genomic AR occupancy was visualized and compared using the Integrated Genome Viewer.

### Immunohistochemistry

Immunohistochemistry staining was performed using the Discovery ULTRA automated stainer from Roche Diagnostics (Indianapolis, IN). In brief, antigen retrieval was performed using a tris/borate/EDTA buffer (Discovery CC1, 06414575001; Roche), pH 8.0 to 8.5, at 95° C for 32 minutes. For AR staining only, 64 minutes of antigen retrieval time was applied. The slides were then incubated with primary antibodies for 1 hour at room temperature with the following dilutions: TMPRSS2 (ab92323), 1:3000; TMPRSS2 (ab214462), 1:200; Androgen Receptor (ab133273), 1:100; ACE2 (R&D Systems, #AF933), 1:400. The antibodies were visualized using the OmniMap anti-Rabbit HRP (05269679001; Roche), and OmniMap anti-Goat HRP (06607233001; Roche) in conjunction with the ChromoMap DAB detection kit (05266645001; Roche). Lastly, the slides were counterstained with hematoxylin and bluing. The specificity of each antibody was first tested on appropriate control tissues before proceeding to staining of the lung sections.

### Gene expression in human lung

The public genomics data repository Gene Expression Omnibus (GEO, ncbi.nlm.nih.gov/geo) was searched for data sets containing expression profiling of samples from non-cancerous human lung and bronchial/airway epithelial cells with samples identified by gender and/or smoking status of subjects. The following data sets were identified: GSE994 (airway epithelial cells from current/former/never smokers), GSE4115 (histologically normal bronchial epithelial cells from smokers with and without lung cancer), GSE7895 (airway epithelial cells from current/former/never smokers), GSE16008 (bronchial epithelial cells from healthy current and never smokers), GSE18385 (large and small airway epithelial cells from healthy current and never smokers), GSE37147 (bronchial epithelial cells from current and former smokers with and without COPD), GSE43696 (bronchial epithelial cells from asthma patients and healthy controls), GSE63127 (small airway epithelial cells from healthy current and never smokers), GSE103174 (lung tissue from smokers and nonsmokers with and without COPD), and GSE123352 (non-involved lung parenchyma from ever and never smokers with lung adenocarcinoma). TMPRSS2, AR, and ACE2 gene expression values were obtained from each data set and analyses for comparisons between groups (for each data set for which gender and/or current smoking status information was available) were performed using R.

### Statistics

For comparisons involving more than two groups, ANOVA with post hoc tests as indicated in figure legends was performed, or Kruskal-Wallis with post hoc tests for IHC results. For comparisons between two groups, t-tests were performed. For correlations between gene expression and smoking pack years, linear models were fit to data. All statistical tests were two-sided. Analyses were performed in R or GraphPad Prism.

### Study approval

Mouse studies were performed under a protocol approved by the Institutional Animal Care and Use Committee (IACUC) of the Cleveland Clinic Lerner Research Institute. Studies using human tissues are deidentified and were deemed to be IRB exempt.

## Acknowledgements

Authors declare no conflicts of interest. This work is supported in part by grants from the Prostate Cancer Foundation and the National Cancer Institute (R01CA172382 and R01CA236780).

